# Cytogenetic comparison of *Cuscuta psorothamnensis* and *C. veatchii* (Convolvulaceae), two species originated from recurrent hybridization between the same diploid parents

**DOI:** 10.1101/2024.04.15.589596

**Authors:** Amalia Ibiapino, Juan Urdampilleta, Miguel A. García, Andrea Pedrosa-Harand, Saša Stefanović, Mihai Costea

**Affiliations:** Instituto Multidisciplinario de Biología Vegetal (IMBIV) - CONICET, Córdoba, Argentina; Real Jardín Botánico-CSIC, Madrid, Spain; Universidade Federal de Pernambuco, Recife, Brazil; Department of Biology, University of Toronto Mississauga, Mississauga, Canada; Department of Biology, University of Wilfrid Laurier, Waterloo, Canada

**Keywords:** *Cuscuta*, allopolyploidy, GISH, parasitic plants, reticulate evolution, speciation

## Abstract

Genus *Cuscuta* L. (Convolvulaceae) exhibits cases of hybridization and allopolyploidy. Section *Denticulatae*, subg. *Grammica*, includes four species: the allopolyploids, *C. veatchii* and *C. psorothamnensis* (2*n* = 60), which originated from two independent reticulation events between the diploids, *C. denticulata* and *C. nevadensis* (2*n* = 30). The allopolyploids are morphologically similar, but are differing in their geographical distribution and host specificity. While cytogenetic data have been reported for *C. veatchii*, this study aims to provide a comparative analysis with *C. psorothamnensis*. To characterize the chromosomal complement of *C. psorothamnensis* and compare it with *C. veatchii*, we used CMA/DAPI banding, FISH, and GISH. The karyotypes of both species displayed similarity in chromosome number, size, and symmetry, and interphase nucleus organization. Both species exhibited a pair of 5S and 35S rDNA sites adjacent on the same chromosome. The number of 5S rDNA sites in *C. psorothamnensis* is variable, with some individuals displaying four, five, and six sites. Our results show: 1) the chromosomal pair carrying adjacent 5S and 35S rDNA in *C. denticulata* is retained in the polyploids; 2) the loss of *C. nevadensis* rDNA sites occurred in both tetraploids; 3) *C. psorothamnensis* and *C. veatchii* are allopolyploids part of a species complex, originated from successive independent hybridization events between *C. denticulata* and *C. nevadensis*; and 4) *C. psorothamnensis* is probably more recent in origin than *C. veatchii* based on the degree of diploidization. This cytogenetic comparison allows us to understand the processes involved in the emergence of new polyploid species by hybridization.

**Main Conclusion:** *Cuscuta psorothamnensis* and *C. veatchii* form a complex of allopolyploid species originating from independent, successive hybridization events between *C. denticulata* and *C. nevadensis*.

## Introduction

Ancestral hybridization is a major evolutionary force, playing a prominent role in the speciation mechanisms of angiosperms (e.g., Stebbins 1958; Vriesendorp and Bakker, 2005; Stull et al. 2023). Reticulation may establish reproductive barriers and contribute to the emergence of characteristics that favor evolutionary success (Soltis and Soltis, 2009, Goulet et al. 2017). Many hybrids exhibit accelerated growth and robust reproductive rate, which provide an advantage to colonizing new habitats, and increase their likelihood of successful establishment (Goulet et al. 2017, Carvalho-Madrigal and Sanín, 2024). Allopolyploids are a common outcome of hybridization and may arise through the fusion of two unreduced gametes (Goulet et al. 2017). Subsequent to hybridization, discernible phenotypic changes may become apparent, characterized by intermediate morphological traits between the parental species (Albuquerque-Lima et al. 2024). In parallel, at the genomic level, processes such as genome homogenization or concerted evolution may occur, leading to the deletion of excess genetic copies or the fixation of genes with adaptive significance (Goulet et al. 2017; Borowska-Zuchowska et al. 2022). Hybridization events can also generate genetic flow between the parental populations, because, if the hybrid is fertile, it can backcross with one or both parents, resulting in introgression. This process can further contribute to the adaptive success of hybrids, reinforcing characteristics such as accelerated growth and robust reproductive rate, explaining their capacity for successful colonization of new environments (Goulet et al. 2017; Carvalho-Madrigal and Sanín, 2024).

Hybridization events lead to genomic changes that may involve the expansion of different classes of transposable elements, followed by loss of genome sequences. Some types of post-hybridization genomic changes are more commonly observed in natural hybrids, such as the loss of rDNA sites through the dosage compensation mechanism that can occur through unequal crossover, which causes the ribosomal cistron of one of the parents to be gradually eliminated (Anamthawat-Jonsoon 2001; Volkov et al. 2017; Li et al. 2021). The chromosomal-level detection of such losses can be accomplished through the application of fluorescence *in situ* hybridization (FISH) and genomic *in situ* hybridization (GISH) techniques (Jiang, 2019). GISH aids in identifying distinct chromosomal sets inherited from each parent involved in the hybrid origin of a species, thereby contributing to research on changes in ploidy levels and potential rearrangements (Silva and Souza 2013; Makonen and Ali, 2023).

*Cuscuta* (dodders) comprises ca. 200 parasitic species with subcosmopolitan distribution (Yuncker 1932; Costea et al. 2015a) and great ecological (Press and Phoenix 2005) as well as economic significance (Lanini and Kogan 2005; Costea and Tardif 2006). The genus shows a remarkable cytogenetic variation, harboring both holocentric or monocentric chromosomes, karyotypes that vary from symmetrical to bimodal, as well as huge variations in the genome size (García and Castroviejo 2003; García et al. 2019; Ibiapino et al. 2022). This wide spectrum of cytogenetic traits within *Cuscuta* contributes to its adaptability and evolutionary success in diverse environments.

*Cuscuta* is currently classified into four subgenera (García et al. 2014; Costea et al. 2015a), with subg. *Grammica* comprising ca. 75% of the species diversity (Stefanović et al. 2007). In addition to diploids with 2*n* = 30, subg. *Grammica* has polyploids, mostly with 2*n* = 60, as well as the highest chromosome counts in the genus, for example 2*n* = 90 in *C. vandevenderi* Costea & Stefanović and 2*n* = 150 in *C. sandwichiana* Choisy (García et al. 2019; Ibiapino et al. 2022). Accompanying this high variation in chromosome numbers, at least 14 cases of independent interspecific hybridization events have been reported in subg. *Grammica* (Stefanović and Costea 2009; Costea and Stefanović2010; Costea et al. 2015b; García et al. 2014).

Sect. *Denticulatae* Yunck. of subg. *Grammica* (Lour.) Peter includes four desert-growing species: *C. denticulata* Engelm., *C. nevadensis* I.M. Johnst. and *C. psorothamnensis* Stefanović, M.A. García & Costea distributed in Western U.S.A., and *C. veatchii* Brandegee found in central Baja California, Mexico (García et al. 2018). This clade is well defined morphologically by the spherically enlarged radicular end of the embryo (Yuncker 1932; Costea et al. 2005), which was suggested to be an adaptation to vivipary and seed germination in the desert (Costea et al. 2005; Olszewski et al. 2020). *Cuscuta denticulata* and *C. nevadensis* are diploid species with a chromosome count of 2*n* = 30, whereas *C. psorothamnensis* and *C. veatchii* are allopolyploids exhibiting a chromosome count of 2*n* = 60 (García et al. 2018). Prior molecular phylogenetic studies of this group (Stefanović and Costea 2008; García et al. 2018) had revealed two cases of strongly supported topological incongruences between plastid- and nuclear-derived phylogenetic trees. These incongruences correspond to the two allopolyploid species, strongly suggesting their hybrid origin from the diploid progenitors. *Cuscuta veatchii* and *C. psorothamnensis* are similar morphologically, differing mainly in their geographical distribution and host specificity (García et al. 2018).

Previous cytogenetic and molecular analyses of *C. veatchii* revealed that none or very few nrITS polymorphisms were present, indicating a complete homogenization by concerted evolution to the rDNA type of *C. nevadensis* (García et al. 2018; Ibiapino et al. 2019). However, the cytogenetic profile of *C. psorothamnensis* has remained unexplored. Given that this group of species is an ideal model for studying speciation through recurrent reticulation accompanied by allopolyploidy, we continue the previous investigations with the following objectives: 1) to conduct a thorough cytogenetic characterization of *C. psorothamnensis*; 2) to compare the cytogenetic profile of this species with the existing characterization of *C. veatchii* to assess their cytogenetic similarity; and 3) to provide a comprehensive discussion on hybrid speciation within section *Denticulatae*. We anticipate that the two hybrid species, *C. psorothamnensis* and *C. veatchii*, share a substantial degree of cytogenetic similarity, while potentially displaying nuanced differences attributable to their likely separate origins and different ages. This study contributes to the broader understanding of hybridization-driven speciation and sheds light on the intricate evolutionary dynamics of dodders.

## Materials and Methods Sampling and material analyzed

Seeds of *C. psorothamnensis* were collected from natural populations in Southern California where the species is endemic (Table 1). After scarification with concentrated sulfuric acid for 20–30 s, seeds were rinsed several times with distilled water, and placed on wet filter paper in Petri dishes to germinate.

**Table 1.** Chromosomal number and number of rDNA sites of *C. psorothamnensis*. Vouchers are deposited at the herbaria of the University of Toronto Mississauga (TRTE) and Wilfrid Laurier University (WLU), Ontario, Canada.

| Species voucher | Locality | 2n | 5S + 35S<br>(total<br>number) | 5S + 35S<br>(adjacent<br>sites only) |
| --- | --- | --- | --- | --- |
| *C. psorothamnensis* Stefanović, M.A. García & Costea<br>*Stefanović SS-16-11* | California; Imperial Co., Hwy 98, E of<br>Imperial Hwy, Ocotillo, 32° 43'29"N 115°<br>59'07"W, 2016 | 60 | 6 + 4 | 2 |
| *Stefanović SS-16-12* | California; San Diego Co., Hwy S2, mi 53,<br>6 mi N of Ocotillo, 32° 47'52"N 116°<br>06'43"W, 2016 | 60 | 4 or 5 and<br>6 in some<br>cells + 4 | 2 |
| *Stefanović SS-16-13* | California; San Diego Co., Hwy S2, mi 50,<br>Carrizo Badlands Overlook, 32°49'43"N<br>116°10'06"W, 2016 | 60 | 4 or 5 and<br>6 in some<br>cells + 4 | 2 |
| *Stefanović SS-16-14* | California; San Diego Co., Hwy S2, mi 47,<br>Mountain Palms Rd., 32°51'55"N<br>116°12'34"W, 2016 | 60 | 6 + 4 | 2 |

### Slide preparation, CMA/DAPI banding, FISH and GISH

Slide preparation was performed using the young shoot tips of seedlings. The material was pretreated with 8-hydroxyquinoline for 24 h at 10 °C, fixed in 3:1 (v/v) ethanol: acetic acid for 2–24h at room temperature, and stored at -20 °C. Subsequently, the material was washed in distilled water, digested in the enzyme Pectinex (Novozimes), and squashed in 60% acetic acid. Double CMA/DAPI staining was performed as described in Ibiapino et al. (2022). The images were captured with a COHU CCD camera attached to a Leica DMLB fluorescence microscope equipped with Leica QFISH software. After image capture, slides were destained for 30 min in Carnoy and 1h in absolute ethanol and stored at -20°C. The destained slides were subjected to FISH according to the protocol detailed by Pedrosa et al. (2002). Two rDNA probes were used. For the 5S rDNA, the four pre-labelled oligos - PLOPs described by Warminal et al. (2018) were ordered from Macrogen end-labelled with Cy3. For the 35S, the pTa71 from wheat (25-28S, 5.8S, and 18S rDNA; Gerlach and Bedbrook, 1979) was labeled by nick translation with Alexa-dUTP (Thermo Scientific).

For GISH, extractions of genomic DNA from *C. denticulata* and *C. nevadensis* were done according to the protocol of Doyle and Doyle (1987). The probes were labeled by nick translation with Cy3-dUTP (*C. denticulata*) or Alexa-dUTP (*C. nevadensis*). The two probes were used at the same time in the hybridization mixture following the same protocol described for fluorescent *in situ* hybridization (FISH). FISH and GISH pictures were obtained as previously described. The selected metaphases were used for chromosome measurement made in Adobe PhotoShop software version 22.3.0.

## Results

All four *C. psorothamnensis* accessions analyzed exhibited 2*n* = 60 chromosomes. The cells selected to be presented in this work had its chromosomes measured and then the values found were averaged. Within these karyotypes, the chromosome size varied by a factor of 2.65, ranging from the smallest (1.44 μm) to the largest pair (3.83 μm), resulting in a total haploid chromosome length of 71.78 μm. The karyotypes displayed a consistent pattern across individuals, with two distinct sets, each consisting of 30 chromosomes. One set consisted of smaller chromosomes with CMA^+^/DAPI^-^ bands in the pericentromeric regions, while the other set comprised larger chromosomes with CMA^0^/DAPI^+^ bands in the pericentromeric region (Fig. 1a, d, and g). The interphase nuclei showed dispersed chromatin with well-defined chromocenters scattered throughout the nucleus (Supplementary Fig. 1g, h and i).

**Fig 1.**
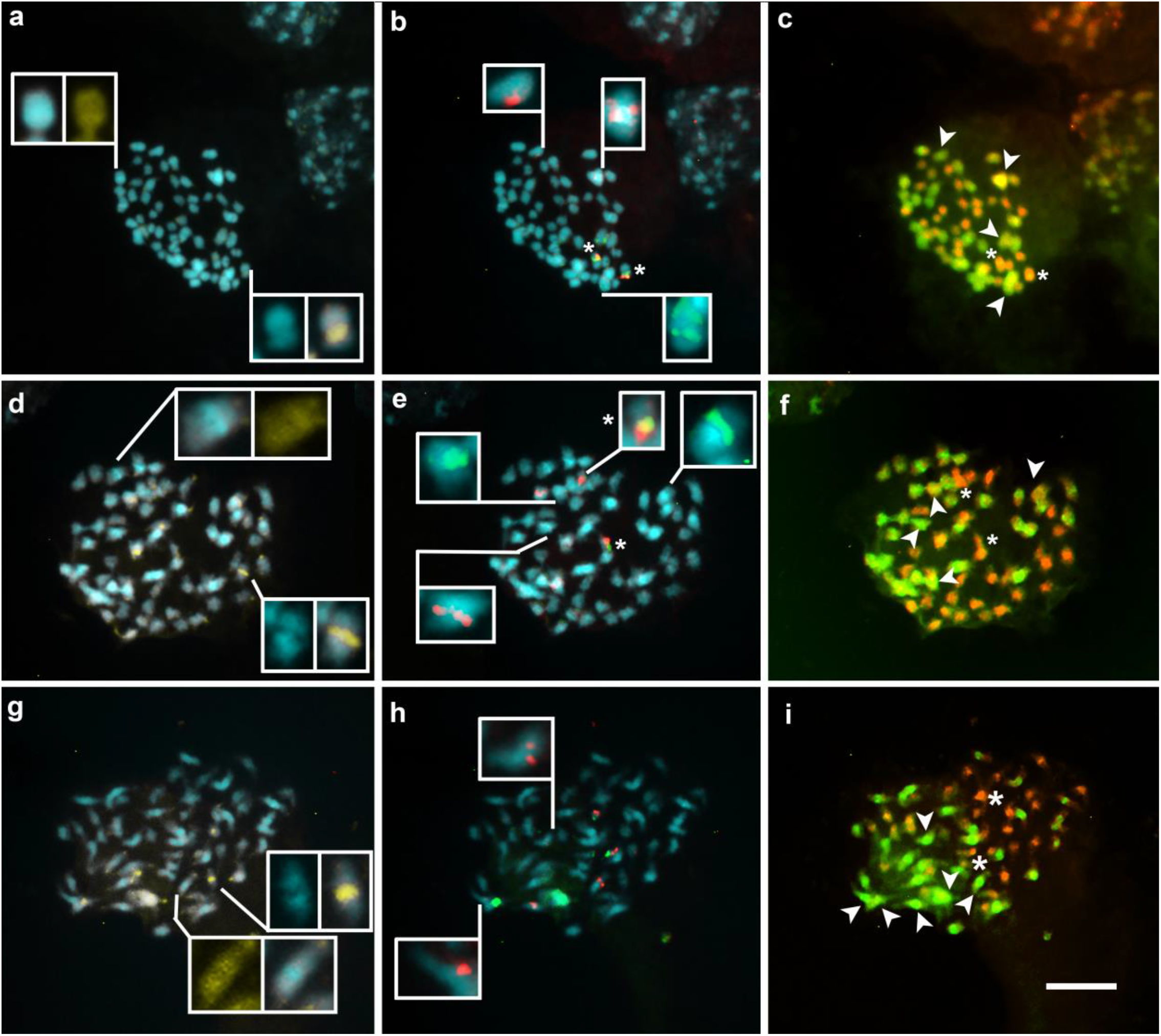
Metaphases from different accessions of *C. psorothamnensis* (a-c SS-16-11 accession and d-i SS-16-13 accession) stained with CMA (yellow) and DAPI (blue) in **a, d** and **g**, FISH using 5S (red) and 35S (green) rDNA probes in **b, e** and **h** and GISH, using the genomic DNA of the parental *C. denticulata* (red) and *C. nevadensis* (green) as a probe in **c, f** and **i**. It is possible to observe different chromosome sets in *C. psorothamnensis*, some chromosomes have CMA^+^/DAPI^-^ bands in the pericentromeric regions (inserts in **a, d** and **g**). The majority of cells from accession SS-16-11 presented four 5S rDNA sites, while accession SS-16-13 presented cells with five or six 5S rDNA sites. The asterisks in **c, f** and **i** highlight the chromosomal pair with adjacent 5S and 35S rDNA on the same chromosome that were inherited from *C. denticulata*, while all other sites were inherited from *C. nevadensis* and are highlighted with arrowheads. Bar in **i** represents 10μm

Fluorescent *in situ* hybridization (FISH) revealed two strongly labeled pairs of 5S rDNA sites and two pairs of 35S rDNA sites in all four accessions, and, in one chromosome pair, 5S and 35S rDNA were adjacent and colocalized with a secondary constriction (Fig. 1b). Some cells, particularly from accessions SS-16-12 and SS-16-13 (Table 1), exhibited one or two additional sites of 5S rDNA, although these extra sites were weakly labeled (Fig. 1e and h). All 35S rDNA sites colocalized with CMA^+^ bands (Fig. 1b, e, and h).

Genomic in situ hybridization (GISH) revealed 30 chromosomes labeled with genomic DNA from *C. denticulata* and 30 chromosomes labeled with genomic DNA from *C. nevadensis*. Chromosomes stained with genomic DNA from *C. denticulata* exhibited staining mainly in pericentromeric regions, while chromosomes stained with genomic DNA from *C. nevadensis* exhibited a disperse staining in some chromosomes. Both probes showed stronger labeling in the pericentromeric regions of the chromosomes. Specifically, the pair of chromosomes with adjacent 5S and 35S rDNA was labeled with *C. denticulata* genomic DNA, while all other rDNA sites were labeled by *C. nevadensis* genomic DNA.

## Discussion

### Cytogenetic similarity between C. *psorothamnensis* and *C. veatchii*

This study revealed a high degree of cytogenetic similarity between the two allopolyploid hybrid species, C. *psorothamnensis* and *C. veatchii*, both exhibiting a chromosome count of 2*n* = 60. Both species presented two distinct chromosome sets, each resembling one of the parental species. Consistent with previous findings for *C. veatchii* (Ibiapino et al. 2019), *C. psorothamnensis* possessed three pairs of 5S rDNA sites and two pairs of 35S rDNA sites, accompanied by interphase nuclei exhibiting diffuse chromatin and well-defined chromocenters. However, *C. psorothamnensis* displayed a higher variation in the number of 5S rDNA sites within the same accession, with individuals showing four, five, or six sites. The number of 5S rDNA sites is most likely six because the other pair of sites exhibited a weak signal, which may make it difficult to visualize in all cells.

GISH results further underscored the similarity between the two hybrids. Both species demonstrated two distinct chromosome sets: 30 chromosomes hybridizing with genomic DNA from *C. denticulata* and another 30 chromosomes labeled with *C. nevadensis* genomic DNA. No recombinant chromosome was detected, as observed in other allopolyploids, such as from the genus *Tragopogon* (Chester et al. 2012). The maintenance of the pair with adjacent 5S and 35S rDNA sites from *C. denticulata* in hybrid karyotypes was evident, but there was a notable loss of rDNA sites from *C. nevadensis*. These data align with previous molecular findings (García et al. 2018), indicating the concerted evolution toward *C. nevadensis* in *C. veatchii* but not in *C. psorothamnensis*, suggesting a more recent independent hybridization event for the latter. The presence of additional 5S rDNA in *C. psorothamnensis*, not eliminated from *C. nevadensis*, as observed in *C. veatchii*, also argues for its younger origin.

Another compelling piece of evidence supporting the younger origin of *C. psorothamnensis* when compared to *C. veatchii* was demonstrated through GISH. The GISH technique, primarily employed to differentiate chromosomes or subgenomes from progenitors in hybrid species, becomes particularly informative when the hybrid is at a more advanced age (Silva and Souza 2013; Ranzam et al. 2017). Genomic probes more strongly labbeled heterochromatic regions of chromosomes. Over time, hybrids can accumulate sequence changes in their genomes or undergo homogenization processes, make emerge some difficult to identify the different subgenomes in their karyotype. When it occur, the use of blocking DNA becomes essential to separate the chromosomal sets effectively (Makonen and Ali 2023). In the case of *C. veatchii*, blocking DNA was essential to obtain a clear-cut result (Ibiapino et al. 2019). In contrast, in *C. psorothamnensis*, a hybrid of more recent origin, using the probes simultaneously yielded clear results, facilitating the distinction of the two chromosome sets.

Hybrids often manifest intermediate characteristics between parental species at both chromosomal and vegetative and reproductive characters levels (e.g., Liu et al. 2009; McKain et al. 2012; Goulet et al. 2017). *Cuscuta psorothamnensis* exemplified this trend by displaying two distinct chromosome sets, one featuring smaller chromosomes with CMA^+^/DAPI^-^ bands in pericentromeric regions, reminiscent of those reported in *C. denticulata*, while the other set presents larger chromosomes, exhibited DAPI^+^/CMA^0^ bands in the pericentromeric region as observed in *C. nevadensis* (Ibiapino et al. 2019; Fig. 2). The interphase nuclei of *C. psorothamnensis* also showed intermediate characteristics. In *C. denticulata*, nuclei are characterized by a more uniform chromatin distribution with small, well-distributed chromocenters, while in *C. nevadensis*, they are larger and well-defined (Ibiapino et al. 2019). The hybrid nuclei presented dispersed chromatin and well-defined chromocenters, indicating a blending of parental traits. The haploid karyotype length in *C. psorothamnensis* was measured at 71.78 μm, not so different from that reported in *C. veachii* with 75.20 μm. It representing a 15% reduction compared to the expected total karyotype length based on both parental species (*C. denticulata* with 35.74 μm and *C. nevadensis* with 49.39 μm; Ibiapino et al. 2019). The genome size in *C. veachii* is 1C = 2.85 Gbp (McNeal et al. 2007), considering the haploid karyotype length, the genome size of *C. psorothamnensis* would be around 1C = 2.72 Gbp.

**Fig 2.**
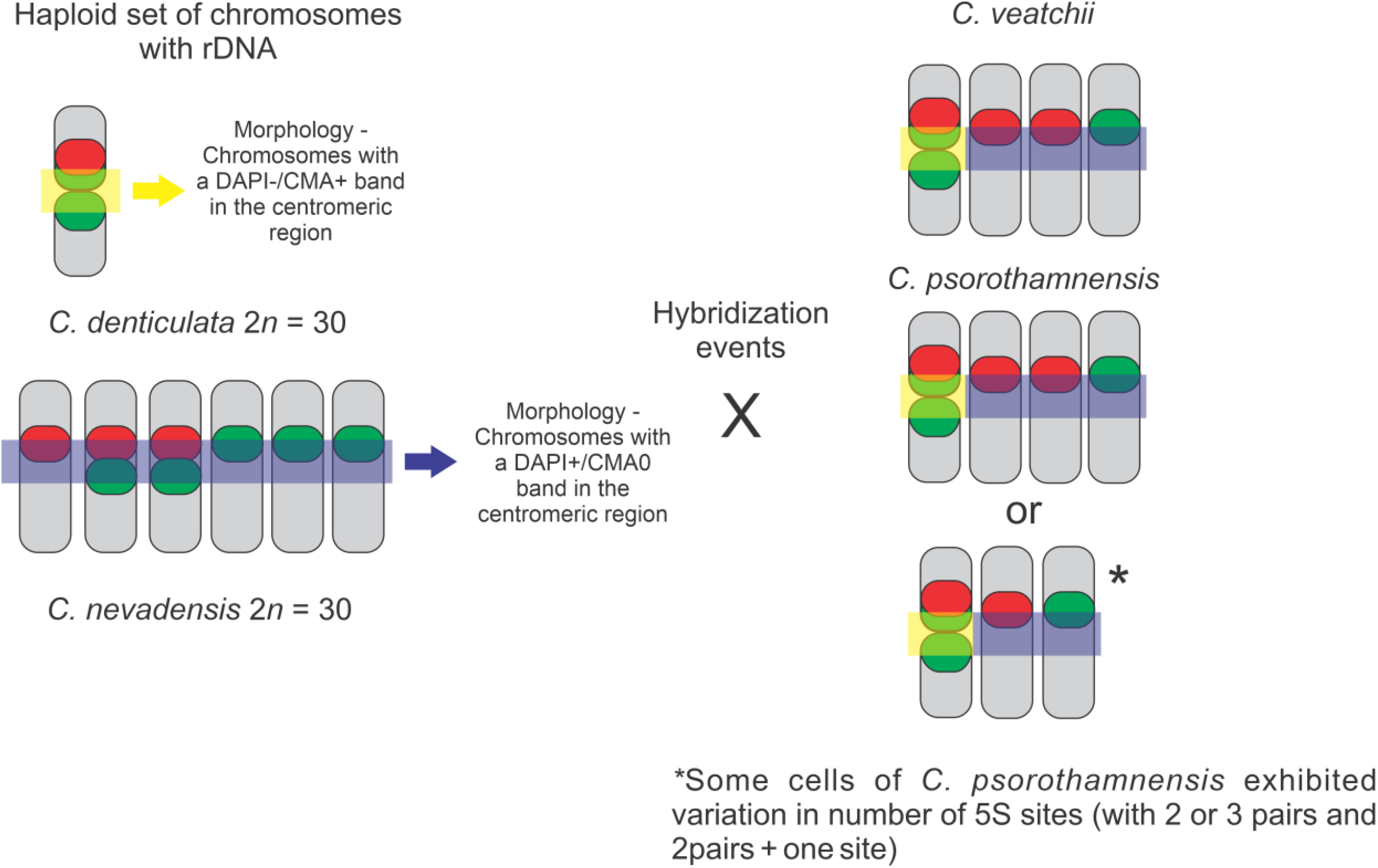
Schematic representation summarizing data previously reported by Ibiapino et al. (2019) combined with the data obtained in the present work. The highlighted yellow and blue shades shows the morphological difference between the chromosomes of *C. denticulata* (with a DAPI^-^ band forming a gap) and *C. nevadensis* (with a DAPI^+^ band and no gap). Both hybrids with *2n* = 60 and two chromosomal sets of different morphology. Only chromosomes containing 5S and 35S rDNA sites were represented in haploid number. The red color represents the 5S rDNA sites, while the green color represents the 35S rDNA sites. A chromosome pair inherited from *C. denticulata* where the 5S and 35S rDNA are adjacent. Those with more rDNA sites inherited from *C. nevadensis*

**Fig 3.**
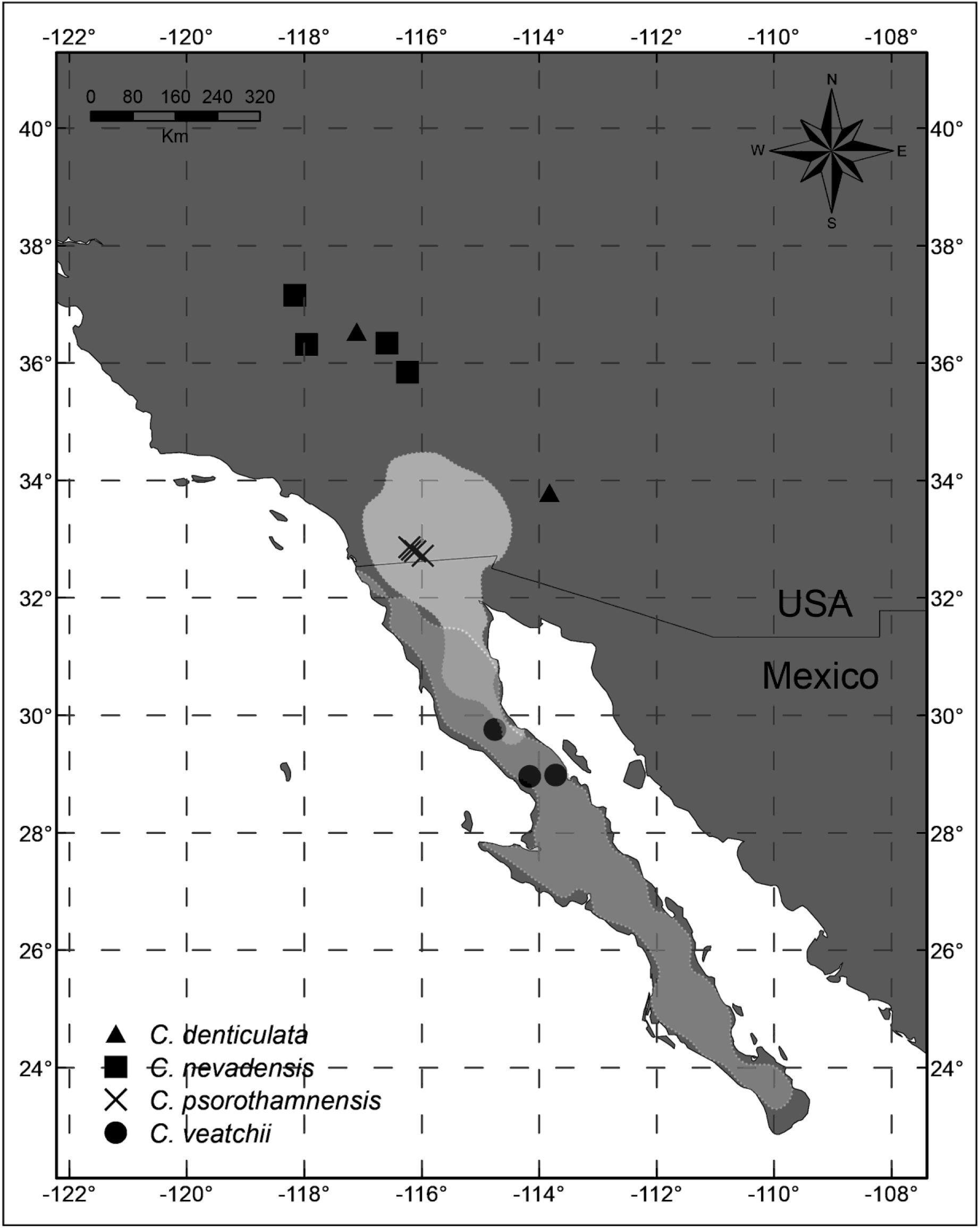
Map showing the geographical position of the accessions analyzed in this work. The shaded areas refer to the regions of occurrence of each of the hosts of the hybrid species, according to Global Biodiversity Information Facility (GBIF – https://www.gbif.org/). The white area refers to *Psorothamnus schottii* and the light gray area to *Pachycormus discolor*. Populations of *C. denticulata* and *C. nevadensis* overlap in their distribution, which would make hybridization events between these two species possible. *Cuscuta veatchii* and *C. psorothamnensis* present distinct ecological characteristics such as host preference and have a different geographic distribution

Hybridization may trigger genome expansions through transposable element (TE) amplification, often due to epigenetic failures like TE methylation (Romero-Soriano et al. 2019; García et al. 2023). Following the initial genome size increase, hybrids typically undergo a subsequent reduction influenced by age, involving the loss of DNA sequences through dosage compensation mechanisms (Kang et al. 2019). In allopolyploid species, a trend toward subgenome dominance emerges, favoring the expression of one parental genome due to extensive duplication post-allopolyploidy. The diploidization process requires the loss of duplicate copies, a phenomenon further accentuated by dosage compensation mechanisms (Kang et al. 2019; Li et al. 2021). The dosage compensation mechanism possibly contributed to the observed reduction in the total length of the haploid karyotype and elimination of excess rDNA copies in *C. psorothamnensis*. Nevertheless, the number of 5S rDNA sites in *C. psorothamnensis* is unexpected. Both parents possess a chromosomal pair with adjacent 5S and 35S rDNA and these are the only sites present in *C. denticulata*, while *C. nevadensis* has three pairs of 5S rDNA sites and five pairs of 35S rDNA, with two chromosomal pairs having adjacent sites (Fig. 2). Our study reveals a discernible reduction in rDNA sites in *C. psorothamnensis*, as observed in *C. veatchii* (Ibiapino et al. 2019).

Typically, rDNA sites are more conserved in artificial hybrids than in natural hybrids (Lee et al. 2011; Volkov et al. 2017). But the presence of two chromosome pairs with only 5S rDNA sites, instead of the expected one that could be inherited from *C. nevadensis*, indicate additional change in rDNA distribution after allopolyploidy and not only elimination, as observed in *Paphiopedilum* (Orchidaceae; Lan and Albert 2011) for example.

### Cuscuta veatchii vs. C. psorothamnensis, the same or different species?

A comprehensive morphometric study of the four species within this section, analyzing over 30 traits (García et al. 2018), highlighted the distinct morphological differences between the two diploid parents, *C. denticulata* and *C. nevadensis*, while the two hybrids, *C. veatchii* and *C. psorothamnensis*, were morphologically indistinguishable. Despite their similar morphology and shared parental origin, the retention of both nrDNA types suggests that *C. psorothamnensis* likely resulted from a more recent independent hybridization event, where ribosomal array homogenization remained incomplete (García et al. 2018). The two hybrids exhibit distinct host preferences not only from each other but also from the parent species. *Cuscuta veatchii* is restricted in central Baja California, Mexico, parasitizing *Pachycormus discolor* (Benth.) Coville (Anacardiaceae), while *C. psorothamnensis* is limited to Anza-Borrego Desert State Park, CA, USA, and grows on *Psorothamnus schottii* (Thor.) Barneby (Fabaceae) (García et al. 2018). The host specificity of the hybrids contrasts sharply with the broader and partially overlapping host ranges observed in the sympatric parental species, *C. denticulata* and *C. nevadensis* (García et al. 2018), suggesting a significant role for host specialization in the cladogenesis of the two hybrid species. Thus, although *C. veatchii* and *C. psorothamnensis* do not meet the morphological criterion for recognition as distinct species, their separate evolutionary histories, ecological differentiation on different hosts, along with their disjunct geographical distributions, strongly indicate that they are two separate species. This is further supported by our cytogenetic results as discussed above.

*Cuscuta* species exhibit varying degrees of host specificity, ranging from “generalists” to “specialists”, with the latter being less common (Gaertner 1950, Costea and Stefanović2009; Costea et al. 2020). Mechanisms underlying host preference in *Cuscuta* are unknown, but they likely control the capacity of the seedlings to detect hosts during foraging (e.g., Benvenuti et al. 2005; Runyon et al. 2006), as well as modulate the signaling with the host during haustoria initiation, penetration, and establishment of the vascular bridge (Jhu and Sinha 2022). Genomic changes after the hybridization may affect possible mechanisms linked to factors that affect downstream gene transcription, changes of hormone status and haustorium formation (Jhu and Sinha 2022). Host specificity and host range also vary widely among parasitic plants (Parker and Riches 1993; Heide-Jørgesen 2008), with host-race formation and host-shifting serving as significant evolutionary drivers in parasitic plants (e.g., Norton and Carpenter 1998; Thorogood et al. 2008; Schneider et al. 2012). The species of *Cuscuta* sect. *Denticulatae* provides a unique opportunity to study the evolution of different host specificity scenarios alongside reticulate evolution and polyploidy. Deeper investigations into the molecular foundation of host preference, continued monitoring of the hybrid populations, and the integration of advanced genomic approaches could unveil novel aspects of host-parasite evolution in *Cuscuta* and parasitic plants more broadly.

## Author Contribution Statement

AI performed the scientific experiments, data collection, and writing of the manuscript. JU contributed by analyzing the data and reviewing the manuscript. MG edited the manuscript, contributed to the collection and identification of the plant material, and discussions of the data, contributed to the writing and reviewing of the manuscript and general discussions. AP-H designed the experiments, supervised, contributed to the writing and reviewing of the manuscript. SS contributed to the collection and identification of the plant material, writing and reviewing the manuscript. MC edited the manuscript, contributed to the collection and identification of the plant material, and discussions of the data contributed to the writing and reviewing of the manuscript and general discussions. All authors contributed to the article and approved the submitted version.

## Acknowledgments

We thank the Consejo Nacional de Investigaciones Científicas y Técnicas (CONICET) for the financing of the Post-Doc scholarship; the Instituto Multidisciplinario de Biología Vegetal (IMBIV) for the infrastructure to develop the work; the Conselho Nacional de Desenvolvimento Científico e Tecnológico (CNPq – financial code 312694/2021-0); and the Coordenação de Aperfeiçoamento de Pessoal de Nível Superior (CAPES), as well as NSERC of Canada Discovery Grant (financial code 326439) for the financial support for the development of this work.

---

**Fig S1.**
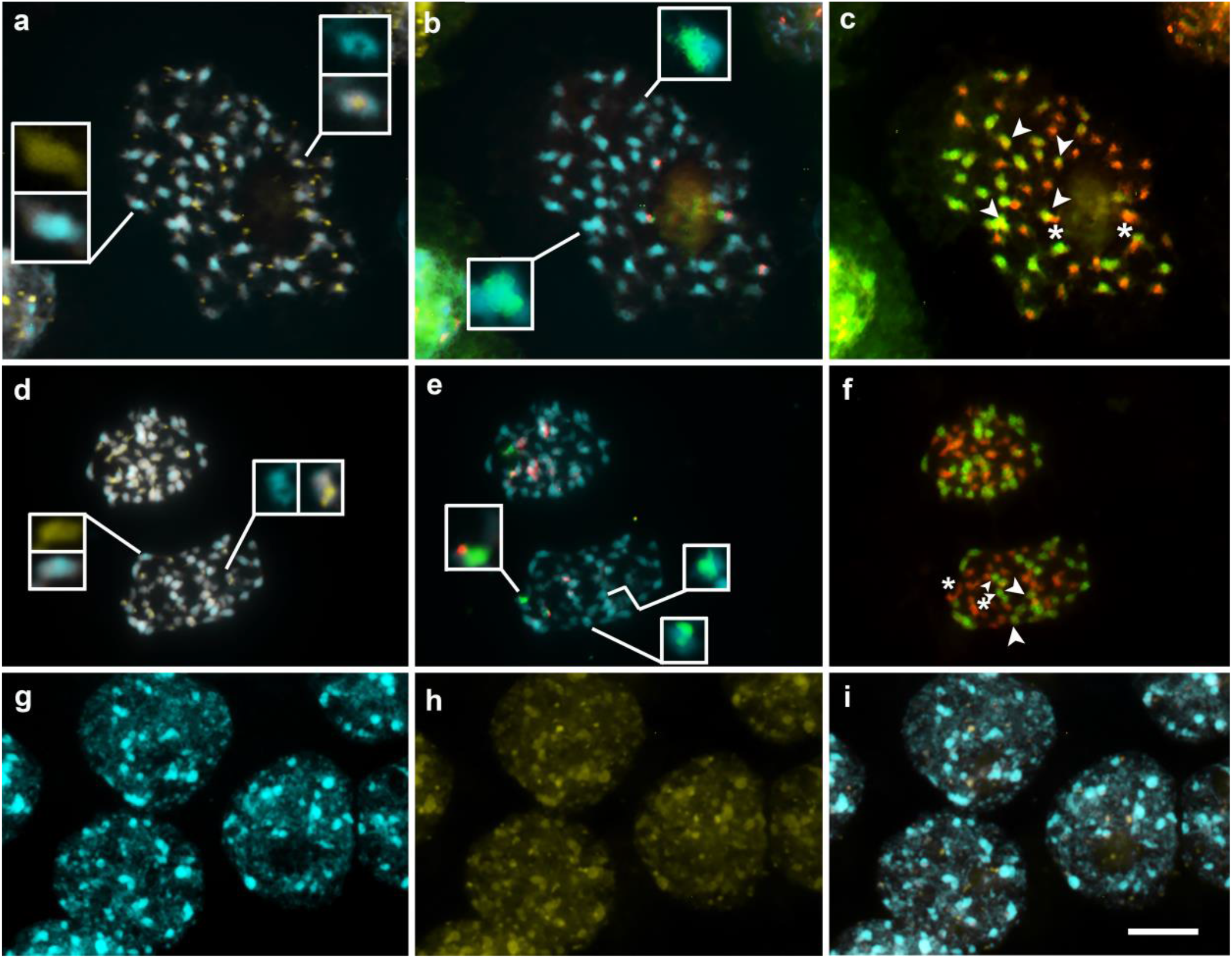
Metaphases from different accessions of *C. psorothamnensis* (a-c SS-16-12 accession and d-i SS-16-14 accession) stained with CMA (yellow) and DAPI (blue) in **a** and **d**, FISH using 5S (red) and 35S (green) rDNA probes in **b** and **e** and GISH, using the genomic DNA of the parental *C. denticulata* (red) and *C. nevadensis* (green) as a probe in **c** and **f**. It is possible to observe different chromosome sets in *C. psorothamnensis*, some chromosomes have DAPI-/CMA+ bands in the pericentromeric regions (inserts in **a** and **d**). In g-i show interphase nuclei of *C. psorothamnensis* stained with DAPI (**g**), CMA (**h**) and the overlap (**i**) showing dispersed chromatin with the presence of small, well-defined chromocenters. The asterisks in **c** and **f** highlight the chromosomal pair with adjacent 5S and 35S rDNA on the same chromosome that were inherited from *C. denticulata*, while all other sites were inherited from *C. nevadensis* and are highlighted with arrowheads. All cells are in the same scale and the bar in **i** represents 10μm

## Notes

### Competing Interest Statement

The authors have declared no competing interest.

## References

Albuquerque-Lima S, Milet-Pinheiro P, Navarro DM, Taylor NP, Zappi DC, Machado IC (2024) Intermediary floral traits between natural hybrid and its parents in the Xiquexique (Cactaceae). Orga Div Evol 10.1007/s13127-023-00634-7.

Anamthawat-Jonsoon (2001) Methods Cell Sci 23: 141–150. 10.1023/A:1013182724179.

Benvenuti S, Dinelli G, Bonetti A, Catizone P (2005) Germination ecology, emergence and host detection in Cuscuta campestris. Weed Res 45: 270–278. 10.1111/j.1365-3180.2005.00460.x.

Borowska-Zuchowska N, Senderowicz M, Trunova D, Kolano B (2022) Tracing the Evolution of the Angiosperm Genome from the Cytogenetic Point of View. Plants (Basel) 11:784. 10.3390/plants11060784.

Carvalho-Madrigal S, Sanín MJ (2024) The role of introgressive hybridization in shaping the geographically isolated gene pools of wax palm populations (genus Ceroxylon). Mol Phyl Evol 193:108013. 10.1016/j.ympev.2024.108013.

Chester M, Gallagher JP, Symonds VV, Silva AVC, Mavrodiev EVM, Leitch AR, Soltis PS, Soltis DE (2012) Extensive chromosomal variation in a recently formed natural allopolyploid species, Tragopogon miscellus (Asteraceae). PNAS 109; 1176–1181. 10.1073/pnas.1112041109.

Costea M, ElMiari H, Farag R, Fleet C, Stefanović S (2020) Cuscuta sect. Californicae (Convolvulaceae) revisited: ‘cryptic’ speciation and host range differentiation. Syst Bot 45; 638–651. 10.1600/036364420X15935294613428.

Costea M, García MA, Stefanović S (2015a) A phylogenetically based infrageneric classification of the parasitic plant genus Cuscuta (Dodders, Convolvulaceae). Syst Bot 40: 269–285. 10.1600/036364415X686567.

Costea M, García MA Baute, K, Stefanović S (2015b). Entangled evolutionary history of Cuscuta pentagona clade: A story involving hybridization and Darwin in the Galapagos. Taxon 64: 1225–1242. 10.12705/646.7.

Costea M, Nesom GL, Tardif FJ (2005) Taxonomic status of Cuscuta nevadensis and C. veatchii (Convolvulaceae) in North America. Brittonia 57: 264–272. 10.1663/0007-196X(2005)057.

Costea M, Stefanović S (2009) Cuscuta jepsonii (Convolvulaceae): An invasive weed or an extinct endemic? Amer J Bot 96: 1744–1750. 10.3732/ajb.0800425.

Costea M, Stefanović S (2010) Evolutionary history and taxonomy of the Cuscuta umbellata complex (Convolvulaceae): Evidence of extensive hybridization from discordant nuclear and plastid phylogenies. Taxon 59: 1783–1800. 10.1002/tax.596011.

Costea M, Tardif FJ (2006) The biology of Canadian weeds. 133. Cuscuta campestris Yuncker, C. gronovii Willd. ex Schult., C. umbrosa Beyr. ex Hook., C. epithymum (L.) L. and C. epilinum Weihe. Canadian J Pl Sci 86: 293–316. 10.4141/P04-077.

Doyle JJ, Doyle JL (1987) A rapid DNA isolation procedure for small quantities of fresh leaf tissue. Phytochem Bull 19:11–15.

Gaertner EE (1950) Studies of seed germination, seed identification and host relationships in dodders. Cuscuta spp. Cornell Univ Agric Exp Stat Mem 294: 1–56.

García S, Kovarik A, Maiwald S, Mann L, Schmidt N, Pascual-Díaz JP, Vitales D, Weber B, Heitkam T (2024) The dynamic interplay between ribosomal DNA and transposable elements: A perspective from genomics and cytogenetics. Mol Biol Evol 2: msae025. doi: 10.1093/molbev/msae025.

García MA, Castroviejo S. (2003) Estudios citotaxonómicos en las especies Ibéricas del género Cuscuta (Convolvulaceae). An Jard Bot Madrid 60:33–44.

García MA, Costea M, Guerra M, García-Ruiz I, Stefanović S (2019) IAPT chromosome data 31. Taxon 68:1374–1380. 10.1002/tax.12176.

García MA, Costea M, Kuzmina M, Stefanović S (2014) Phylogeny, character evolution, and biogeography of Cuscuta (dodders; Convolvulaceae) inferred from coding plastid and nuclear sequences. Am J Bot 101:670–90. 10.3732/ajb.1300449.

García MA, Stefanović S, Weiner C, Olszewski M, Costea M (2018) Cladogenesis and reticulation in Cuscuta sect. Denticulatae (Convolvulaceae). Org Divers Evol 18:383–398. 10.1007/s13127-018-0383-5.

Gerlach WL, Bedbrook JR (1979) Cloning and characterization of ribosomal RNA genes from wheat and barley. Nucleic Acids Res 7:1869–1885. doi: 10.1093/nar/7.7.1869.

Goulet BE, Roda F, Hopkins R (2017) Hybridization in Plants: Old Ideas, New Techniques. Plant Physiol 173:65–78. 10.1104/pp.16.01340.

Heide-Jørgesen H (2008) Parasitic flowering plants. Brill, Leiden, Netherlands.

Ibiapino A, García MA, Amorim B, Báez M, Costea M, Stefanović S, Pedrosa-Harand A (2022) The evolution of cytogenetic traits in Cuscuta (Convolvulaceae), the genus with the most diverse chromosomes in Angiosperms. Front Plant Sci 13:842260. 10.3389/fpls.2022.842260.

Ibiapino A, García MA, Ferraz ME, Costea M, Stefanović S, Guerra M (2019) Allopolyploid origin and genome differentiation of the parasitic species Cuscuta veatchii (Convolvulaceae) revealed by genomic in situ hybridization. Genome 62:467–475 10.1139/gen-2018-0184.

Jhu MY, Sinha NR (2022) Cuscuta species: Model organisms for haustorium development in stem holoparasitic plants. Front Plant Sci 13:1086384. 10.3389/fpls.2022.1086384.

Jiang J (2019) Fluorescence in situ hybridization in plants: recent developments and future applications. Chrom Res 27:153–165. 10.1007/s10577-019-09607-z.

Kang Z, Xiaowu W, Feng C (2019) Plant Polyploidy: Origin, Evolution, and Its Influence on Crop Domestication. Hortic Plant Jour 5:231–239. 10.1016/j.hpj.2019.11.003.

Lanini WT, Kogan M (2005) Biology and management of Cuscuta in crops. Int J Agric Nat Res 32: 127–141.

Lan T, Albert VA (2011) Dynamic distribution patterns of ribosomal DNA and chromosomal evolution in Paphiopedilum, a lady’s slipper orchid. BMC Plant Biol 11: 126. 10.1186/1471-2229-11-126.

Lee YI, Chang FC, Chung MC (2011) Chromosome pairing affinities in interspecific hybrids reflect phylogenetic distances among lady’s slipper orchids (Paphiopedilum). Ann Bot 108:113–21. 10.1093/aob/mcr114.

Li Z, McKibben MTW, Finch GS, Blischak PD, Sutherland BL, Barker MS (2021) Patterns and processes of diploidization in land plants. Ann Rev Pl Biol 72:387–410. 10.1146/annurev-arplant-050718-100344.

Liu J, Wang H, Yu L, Li D, Li M (2009) Morphology and cytology of flower chimeras in hybrids of Brassica carinata × Brassica rapa. African J Biotech 8:801–806. 10.1016/j.sajb.2019.08.002.

McKain MR, Wickett N, Zhang Y, Ayyampalayam S, McCombie WR, Chase MW, Pires JC, dePamphilis CW, Leebens-Mack J (2012) Phylogenomic analysis of transcriptome data elucidates co-occurrence of a paleopolyploid event and the origin of bimodal karyotypes in Agavoideae (Asparagaceae). Am J Bot 99:397–406. 10.3732/ajb.1100537.

McNeal JR, Kuehl JV, Boore JL, Pamphilis CW (2007). Complete plastid genome sequences suggest strong selection for retention of photosynthetic genes in the parasitic plant genus Cuscuta. BMC Plant Biol 7:57. doi: 10.1186/1471-2229-7-57.

Mekonem AA, Ali A (2023) A review on principles of FISH and GISH and its role in cytogenetic study. Glob Res Envirom Sustainab 1:15–26.

Norton DA, Carpenter MA (1998) Mistletoes as parasites: Host specificity and speciation. Trends Ecol Evol 13: 101–105. 10.1016/s0169-5347(97)01243-3.

Olszewski M, Dilliott M, García-Ruiz I, Bendarvandi B, Costea M (2020) Cuscuta seeds: Diversity and evolution, value for systematics/identification and exploration of allometric relationships. PloS ONE 15: 0234627. 10.1371%2Fjournal.pone.0234627.

Parker C, Riches CR (1993) Parasitic weeds of the world. Biology and control. CAB International, Wallingford, UK.

Pedrosa A, Sandal N, Stougaard J, Schweizer D, Bachmair A (2002) Chromosomal map of the model legume Lotus japonicus. Genetics 161:1661–1672. 10.1093/genetics/161.4.1661.

Press MC, Phoenix GK (2005) Impacts of parasitic plants on natural communities. New Phytol 166: 737–751. 10.1111/j.1469-8137.2005.01358.x.

Ramzan F, Younis A, Lim KB (2017) Application of genomic in situ ybridization in horticultural science. Int J Genomics 2017:7561909. 10.1155/2017/7561909.

Romero-Soriano V, Burlet N, Vela D, Fontdevila A, Vieira C, García Guerreiro MP (2016) Drosophila females undergo genome expansion after interspecific hybridization. Genome Biol Evol 8:556–61. 10.1093/gbe/evw024.

Runyon JB, Mescher MC, De Moraes CM (2006) Volatile chemical cues guide host location and host selection by parasitic plants. Science 313: 1964–1967. 10.1126/science.1131371.

Schneider AC, Colwell AEL, Schneeweiss GM, Baldwin BG (2016) Cryptic host-specific diversity among western hemisphere broomrapes (Orobanche s.l., Orobanchaceae). Ann Bot 118: 1101–1111. 10.1093/aob/mcw158.

Silva GS, Souza MM (2013) Genomic in situ hybridization in plants. Genet Mol Res 12:2953–65. 10.4238/2013.August.12.11.

Stebbins GL (1958) On the hybrid origin of the angiosperms. Evolution 12: 267–270.

Stull GW, Pham KK, Soltis PS, Soltis DE (2023) Deep reticulation: the long legacy of hybridization in vascular plant evolution. The Plant Journal 114; 743–766. 10.1111/tpj.16142.

Soltis PS, Soltis DE (2009) The role of hybridization in plant speciation. Annu Rev Plant Biol 60:561–88. 10.1146/annurev.arplant.043008.092039.

Stefanović S, Kuzmina M, Costea M (2007) Delimitation of major lineages within Cuscuta subgenus Grammica (dodders; Convolvulaceae) using plastid and nuclear DNA sequences. Am J Bot 94: 568–589. 10.3732/ajb.94.4.568.

Thorogood CJ, Rumsey FJ, Harris SA, Hiscock SJ (2008) Host-driven divergence in the parasitic plant Orobanche minor Sm. (Orobanchaceae). Mol Ecol 17:4289–4303. 10.1111/j.1365-294x.2008.03915.x.

Volkov RA, Panchuk II, Borisjuk NV, Hosiawa-Baranska M, Maluszynska J, Hemleben V (2017) Evolutional dynamics of 45S and 5S ribosomal DNA in ancient allohexaploid Atropa belladonna. BMC Plant Biology 17:21. 10.1186/s12870-017-0978-6.

Vriesendorp B, Bakker FT (2005) Reconstructing patterns of reticulate evolution in angiosperms: what can we do? Taxon 54: 593–604. 10.2307/25065417.

Yuncker TG (1932) The genus Cuscuta. Mem Torrey Bot Club 18: 113–331.

